# Genomic diversity and antimicrobial resistance of *Haemophilus* colonising the airways of young children with cystic fibrosis

**DOI:** 10.1101/2020.11.23.388074

**Authors:** Stephen C. Watts, Louise M. Judd, Rosemary Carzino, Sarath Ranganathan, Kathryn E. Holt

**Affiliations:** Department of Biochemistry and Molecular Biology, Bio21 Molecular Science and Biotechnology Institute, University of Melbourne, 3010, Australia; Department of Infectious Diseases, Central Clinical School, Monash University, Melbourne, Victoria 3004, Australia; Infection and Immunity, Murdoch Children’s Research Institute; Department of Paediatrics, University of Melbourne, 3010, Australia; London School of Hygiene & Tropical Medicine, London WC1E 7HT, UK

## Abstract

Respiratory infection during childhood is a key risk factor in early cystic fibrosis (CF) lung disease progression. *Haemophilus influenzae* (*Hi*) and *Haemophilus parainfluenzae* (*Hpi*) are routinely isolated from the lungs of children with CF, however little is known about the frequency and characteristics of *Haemophilus* colonisation in this context. Here, we describe detection, antimicrobial resistance (AMR) and genome sequencing of *Hi*/*Hpi* isolated from sputum, cough swab, and bronchoalveolar lavage samples regularly collected from 147 participants aged ≤12 years enrolled in the Australian Respiratory Early Surveillance Team for Cystic Fibrosis (AREST CF) program. The frequency of colonisation per visit was 4.6% for *Hi* and 32.1% for *Hpi*, 80.3% of participants had *Hi* and/or *Hpi* detected on at least one visit, and using genomic data we estimate 15.6% of participants had persistent colonisation with the same strain for at least two consecutive visits. Colonising strains were genetically highly diverse and AMR was common, with 52% of *Hi* and 82% of *Hpi* displaying resistance to at least one drug. The genetic basis for AMR could be identified in most cases; novel determinants include a new plasmid encoding *bla*_TEM-1_ (ampicillin resistance), a new inhibitor-resistant *bla*_TEM_ allele (augmentin resistance), and previously unreported mutations in chromosomally-encoded genes (*pbp3*, ampicillin resistance; *folA* and *folP*, co-trimoxazole resistance; *rpoB*, rifampicin resistance). Acquired AMR genes were significantly more common in *Hpi* than *Hi* (51% vs 21%, p=0.0107) and were mostly associated with the ICE*Hin* mobile element carrying *bla*_TEM-1_, resulting in higher rates of ampicillin resistance in *Hpi* (73% vs 30%, p=0.0004). The genome data identified six potential instances of *Haemophilus* transmission between participants, three of which involved participant pairs who attended the clinic on the same day. The high prevalence of *Haemophilus* colonisation and high burden of antimicrobial use in children with CF likely provides a reservoir for emergence and spread of AMR as well as a source of infections.

## Introduction

Cystic fibrosis (CF) is a common inherited genetic disorder caused by deleterious mutations in the cystic fibrosis transmembrane conductance regulator gene (1). Although the disease is multi-systemic, the primary cause of morbidity and mortality result from pulmonary dysfunction. CF lung disease manifests as delayed mucociliary clearance and mucus adhesion leading to recurrent and chronic bacterial infections (2), which elicit adverse host inflammation response resulting in bronchiectasis and ultimately respiratory failure (3,4). Management of bacterial lung infections is essential in CF disease trajectory and can be controlled through antimicrobial therapy. However, antimicrobial resistance (AMR) is inevitably acquired through various mechanisms and has severe clinical consequences that are highly relevant in an era of antimicrobial custodianship.

There are a small number of bacterial species that predominantly cause CF lung infections, including *Pseudomonas aeruginosa, Burkholderia cepacia* complex, *Staphylococcus aureus, Stenotrophomonas maltophilia*, and *Haemophilus influenzae* (5). Importantly, acute respiratory infection in newborns with CF is an established risk factor for early disease development and progression (6).

*H. influenzae* (*Hi*) and *Haemophilus parainfluenzae* (*Hpi*) are among the most common *Haemophilus* species that colonise the respiratory tract of children in early life (7,8). *Hi* and *Hpi* are both considered opportunistic pathogens and can cause invasive disease, though *Hpi*-related disease is less frequently observed and *Hpi* is recognised as having a lower pathogenic capacity than *Hi*. Both *Hi* and *Hpi* are routinely isolated from the respiratory tract of children with CF, particularly during episodes of disease exacerbation (9). Although many of the classical pathogens involved in CF lung disease have been well studied, less is known about the role of *Haemophilus* species during the critical period of early childhood. Such knowledge is essential as it is increasingly recognised that CF lung disease commences soon after diagnosis in early infancy and progresses thereafter (10,11). Further insight into the epidemiology and resistance profiles of these early colonising and infecting bacteria will inform future treatment practices.

The emergence and accumulation of AMR in *Hi* and *Hpi* is common. Generally, *Hi* and *Hpi* infections are treated with β-lactams such as extended-spectrum penicillins or cephalosporins (12). Other drugs are often used in combination with or as an alternative to β-lactams, and include anti-folates, quinolones, and macrolides. Resistance to these drugs typically arise through either acquisition of horizontally transferred resistance genes or mutations in chromosomally encoded protein targets (12). Acquired AMR genes in *Hi* and *Hpi* are frequently localised within mobile genetic elements such as *ICEHin* or small plasmids (13–15), which appear to have facilitated emergence of multidrug resistant *Hi* and *Hpi* strains in recent years (16,17).

*Hi* is known to produce polysaccharide capsule, which can be classified into six serotypes (Hia through Hif) and is an invasive virulence determinant (18). Strains that do not produce capsule are designated non-typeable *Hi* (NTHi). The introduction of the highly effective Hib conjugate vaccines caused a marked reduction of Hib-related disease incidence but consequently resulted in an increased prevalence of NTHi-related disease (19); NTHi is now more commonly isolated from children with CF than any encapsulated *Hi* serotype (20,21). *Hpi* is generally less well characterised and the role it may have in CF disease is unclear. There is no detailed description of encapsulated *Hpi* though there is increasing evidence that some *Hpi* strains could express polysaccharide capsule (17). Moreover, there is a stark lack of *Hpi* genomic data compared to *Hi* despite it occupying a similar niche. Here we investigate the prevalence, genomic diversity and AMR phenotypes of *Hi* and *Hpi* colonising the airways of children with CF, recruited at the Royal Children’s Hospital (RCH), Melbourne, Australia.

## Methods

### Participant recruitment, sample and data collection

Participants in this study are a subset of those enrolled in the Australian Respiratory Early Surveillance Team for Cystic Fibrosis (AREST CF) birth cohort who meet the following inclusion criteria: diagnosed with CF, under 12 years of age, resident in catchment area, and presented to the RCH CF clinic between February 2016 to February 2017 (10). Respiratory samples (bronchoalveolar lavage (BAL), sputum, or cough swabs) were routinely collected from participants during regular visits and cultured in the RCH microbiological diagnostics lab as described previously (10). During the one-year study period, 847 samples collected from 147 study participants were analysed and yielded 39 isolates identified as *Hi* and 272 identified as *Hpi* (identified using the X and V factor test). Isolates were tested for susceptibility to ampicillin (AMP), augmentin (AMC), cefotaxime (CTX), co-trimoxazole (STX), and rifampicin (RIF) using disc diffusion with CLSI breakpoints.

### Bacterial isolates, sequencing, and assembly

A total of 162 of the 311 *Haemophilus* isolates (n=30, 77% of those biochemically identified as *Hi* and n=132, 48.5% as *Hpi*) were successfully resuscitated, subcultured and transferred to the University of Melbourne for whole genome sequencing (WGS). Isolates were plated on chocolate agar and incubated at 37°C in microaerophilic conditions for 48 hours. Colonies were harvested and DNA extracted using GenFindV2 (Beckman Coulter), using proteinase K for bacterial lysis according to manufacturer’s instructions. Short-read DNA libraries were prepared for all isolates with Nextera XT (Illumina) and subsequently sequenced on the Illumina MiniSeq platform, generating paired-end reads of 151 bp each. DNA samples for long-read sequencing were prepared for a subset of 14 isolates using GenFindV2 (Beckman Coulter), a barcoded ligation library was prepared (SQK-LSK108, EXP-NBD103), and sequenced via Oxford Nanopore MinION on a R9.4.1 flow cell.

A total of 107 isolates (24 *Hi*, 83 *Hpi*) were successfully sequenced via Illumina and passed quality control, each yielding ≥150,000 high-quality reads (**Figure S1** and **File S1**). Centrifuge v1.0.4b (22) was used to categorise isolates as either: (a) pure *Hi* or *Hpi*, defined as one of these species at ≥50% relative abundance and the next most common species <20% relative abundance; (b) contaminated *Hi* or *Hpi*, defined as one of these species at ≥50% relative abundance and a second species also highly represented (≥20% relative abundance); or (c) other, where neither of these species exceeded 50% relative abundance (**Figure S1**). Strain multiplicity for pure *Hi* and *Hpi* cultures was assessed by comparing the ratio of heterozygous to homozygous single nucleotide variant (SNV) calls (methods below) against an empirically determined threshold (*Hi*, ≥0.025 or *Hpi*, ≥0.100, calculated from public datasets); samples exceeding this threshold were considered to be mixed cultures and were excluded from further analysis (**Figure S1**). Genomes were assembled with Unicycler v0.4.7 (23), using Illumina data in all cases and complemented by MinION data where available. Read data were deposited under the NCBI BioProject accession PRJNA668428 (see **File S1** for individual accessions) and assemblies were deposited as a FigShare dataset (https://doi.org/10.26180/13251419).

### Population structure analysis

The *Hi* and *Hpi* short-read Illumina data generated in this study (n=107), and publicly available read sets for previously sequenced genomes of these species (n=891; summarised in **Table S1**), were subjected to SNV detection, phylogenetic and population structure analyses. SNVs were called using Bowtie2 v2.2.9 (for read mapping) and SAMtools v1.9 (for variant calling) via the RedDog pipeline v1beta.11 (https://github.com/katholt/reddog), using *Hi* strain Rd KW20 (accession GCA_000027305.1) and *Hpi* strain T3T1 (accession GCA_000210895.1) as reference genomes. For each species, maximum-likelihood core-genome SNV phylogenies were inferred from alignments of SNV alleles at sites present in ≥95% of genomes (264,940 variable sites for 901 *Hi* genomes, 329,046 variable sites for 97 *Hpi* genomes), using IQ-TREE v2.1.0 (24) and visualised with ggtree v1.14.6 (25) in R v3.5.2 (26). *Hi* capsular serotype loci were detected from genome assemblies using hicap v1.0.0 (27), and sequence types (STs) were assigned to *Hi* read sets using SRST2 v.0.2.0 (28) with the *Hi* multi-locus sequence typing (MLST) database (29).

Discriminant Analysis of Principal Components (DAPC) (30) was conducted to explore the relationship between bacterial population structure and sample source using *k*-mers (of length *k*=16) extracted from assemblies. Frequencies of *k*-mers were counted in each assembly with fsm-lite v1.0 (https://github.com/nvalimak/fsm-lite), and a presence-absence matrix constructed. Due to memory limitations, random sets of 500,000 *k*-mers were selected from the presence-absence matrix to use as input for DAPC, which was performed with the R package adegenet v2.1.1 (31) in triplicate using different random *k*-mer subsets to ensure stability of results (and additionally using the core-genome SNVs called from reads).

### Analysis of antimicrobial determinants

RCH isolates were investigated for known and novel AMR determinants. Reads and assemblies were screened using SRST2 v0.2.0 and BLAST v2.7.1, respectively, to identify alleles of horizontally transferred AMR genes curated in the ARG-ANNOT database (32). Exact matches for translated *bla*_TEM_ gene sequences were identified in NCBI AMR database with BLAST to infer spectrum of activity and inhibitor resistance.

Mutations in chromosomally encoded antimicrobial target genes (*ftsI, folA, rpoB*, and *pbp* genes) (33–43) were also investigated. An exhaustive search for PBP genes present in the *Hi* and *Hpi* reference genomes was performed by aligning translated gene sequences to all curated PBP protein sequences available in the Swiss-Prot database (44) to identify those with ≥80% coverage and ≥70% identity (**Table S2**). Nucleotide sequences for target genes *(ftsI, folA, rpoB*, and *pbp* genes) were extracted from RCH isolate assemblies, and the translated amino acid sequences aligned using MAFFT v7.407 (45). Each alignment position was compared to the consensus sequence for all isolates that were sensitive to the relevant antimicrobial. Positions that varied were tested for statistical association with the corresponding antimicrobial susceptibility phenotype (expressed as a binary variable, insensitive (I/R) vs sensitive (S)) using Fisher’s exact test, and using linear mixed models (LMM) to correct for population structure by including a genetic relatedness matrix calculated from the alignment of biallelic SNVs. LMMs were fitted with GEMMA v0.98.1 (46) and significance was assessed by the Wald test. The resulting *p*-values were adjusted for multiple testing using Benjamini-Hochberg correction on a per-gene basis. Significant variants (*p*-value < 0.05) are reported in **Table S4-S5** and distribution of variants in isolates are detailed in **File S2**.

### Identification of persistent and transmitted strains

“Strains” were defined as groups of closely related isolates, and identified initially using complete-linkage hierarchical clustering based on SNV distances derived from the global core-genome alignment (**Figure S2**). To capture isolate pairs with inflated SNV counts due to small numbers of recombination events, the SNV distance thresholds for strain definition were set to ≤2,000 for *Hi* and ≤4,000 for *Hpi* (**Figure S2A and C**). Precise pairwise SNV distances within each potential strain group were obtained by mapping all isolates to the best within-group genome assembly (lowest N50) using RedDog v1beta.11 as described above. Recombination blocks were identified by comparing pairwise SNV densities within discrete 4-kbp windows along the genome with mean pairwise SNV count for all windows, using a binomial test and Bonferroni correction to account for multiple testing within each strain group (**Figure S3**). Genetic distance between isolate pairs was then defined as non-recombinant SNVs plus the number of recombinant blocks; and strains defined as groups of isolates with pairwise genetic distance ≤20 (**Figure S2B and D**).

## Results

### Detection and sequencing of *Haemophilus* isolates

During the one-year study period, 147 AREST CF study participants receiving treatment at the RCH CF specialist clinic were screened for presence of *Hi* or *Hpi* in the respiratory tract during regular clinic visits and during hospitalisation of pulmonary exacerbation. Participant characteristics are given in **Table 1**. Thirty (20.4%) participants tested culture-positive for colonisation with *Hi* in ≥1 sample, and 111 (75.5%) participants for *Hpi*. Only 29 participants (19.7%) had no *Haemophilus*-positive samples; these individuals did not differ by age or gender but contributed fewer samples each (mean 4.6 vs 6.0, p=0.012 using two-sample Kolmogorov-Smirnov test). The overall frequency of colonisation per visit was 4.6% for *Hi* and 32.1% for *Hpi*, with 86 participants (58.5%) presenting with either *Hi* or *Hpi* on two or more occasions. Several participants had both *Hi* and *Hpi* detected during the same (n=10, 6.8%) or different (n=15, 10.2%) visits.

**Table 1.** Study participant characteristics.

|  | **All participants**<br>**(Lab reported species ID)** | **Participants with WGS**<br>**data (WGS species ID)** |
| --- | --- | --- |
| **All participants** |  |  |
| Total participants, count | 147 | 59 |
| Female, count (%) | 64 (43.5%) | 23 (38.9%) |
| Mean age at first sample, years (range) | 5.7 (0.08-11.8) | 4.1 (0.10-8.9) |
| Mean number of samples, count (range) | 5.8 (1-12) | 6.2 (2-12) |
| ≥1 *Hi* positive sample, count (%) | 30 (20.4%) | 21 (35.6%) |
| ≥1 *Hpi* positive sample, count (%) | 111 (75.5%) | 55 (93.2%) |
| Zero *Hi* or *Hpi* positive samples, count (%) | 29 (19.7%) | - |
| **Participants with ≥1 *Haemophilus* culture-positive sample** |  |  |
| Total participants, count | 118 | 59 |
| Female, count (%) | 54 (45.8%) | 23 (39.0%) |
| Mean age at first sample, years (range) | 5.3 (0.08-11.8) | 4.1 (0.10-8.9) |
| Mean number of samples, count (range) | 6.0 (2-12) | 6.2 (2-12) |
| Mean number of *Hi* positive samples, count (range) | 0.3 (0-3) | 0.5 (0-3) |
| Mean number of *Hpi* positive samples, count (range) | 2.3 (0-7) | 2.8 (0-7) |

A subset of 162 *Haemophilus* isolates from 59 participants (50% of culture-positive individuals, representative in terms of age and gender, see **Table 1**) were subjected to WGS, of which 89.5% were sequenced successfully. WGS data revealed some species misidentifications (5.5%) and mixed cultures (13%), leaving 24 *Hi* and 83 *Hpi* isolates for further analysis (see **Figure S1** and **Methods**).

### Genetic diversity and population structure

The *Hi* isolates collected in this study were highly diverse (median 2.3% nucleotide divergence in core genome), and comparison with publicly available WGS data from other studies (n=877 genomes, see **File S1**) indicates RCH CF isolates are distributed across the global species phylogeny (red tips in **Figure 1A**). All but two RCH CF *Hi* genomes had no detectable capsule biosynthesis (*cap*) locus and fell outside the small number of lineages typically associated with encapsulation (27) (coloured branches in **Figure 1A**), and are thus predicted to be unencapsulated or non-typeable. Two RCH CF isolates (drug susceptible, sequence type (ST) 18, from the same patient) carried intact copies of the *cap-e* locus and fell within the lineage typically associated with serotype e (green branches in **Figure 1A**), thus are predicted to express serotype e capsules.

**Figure 1.**
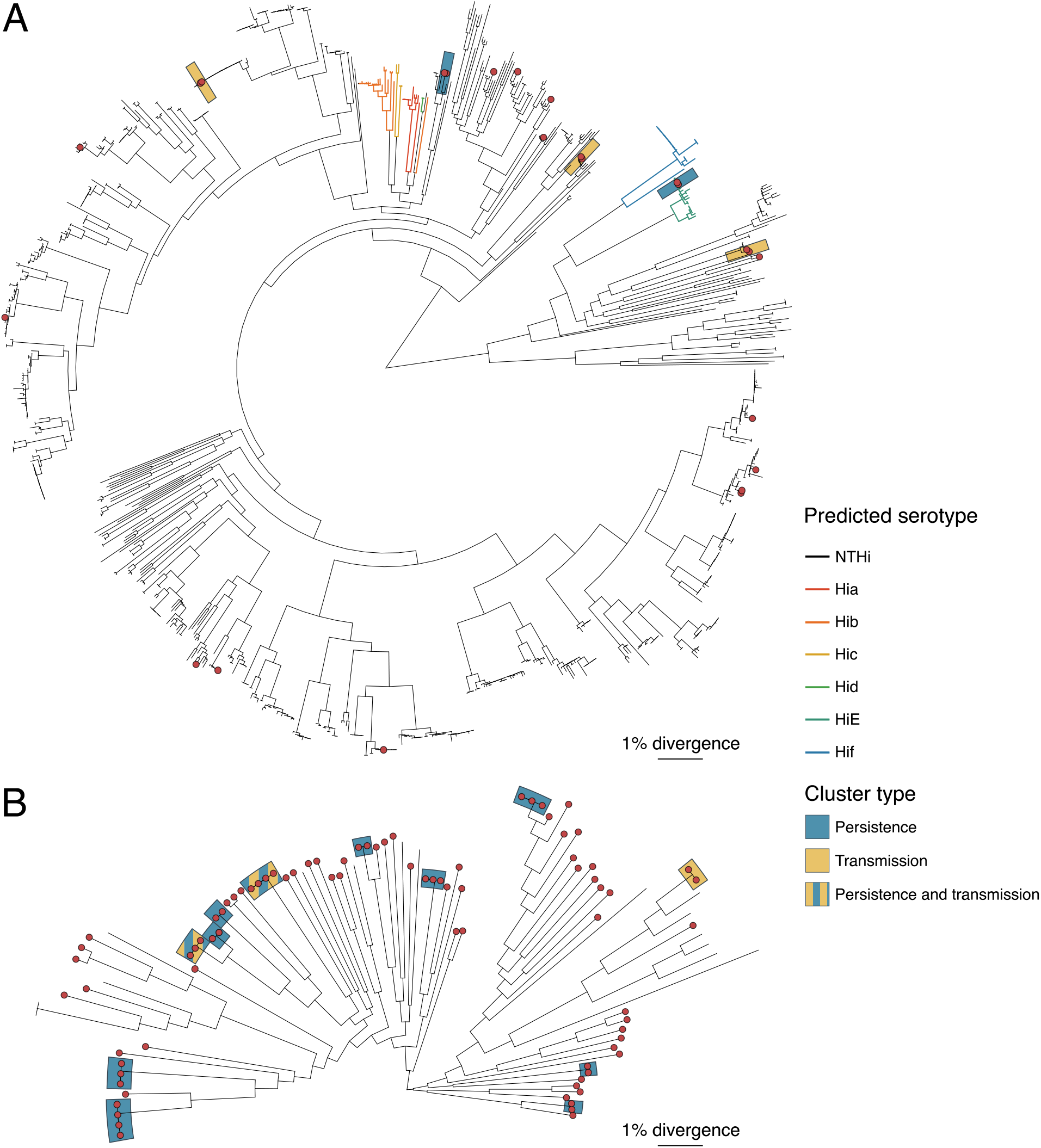
Maximum-likelihood phylogenies for (A) *Hi* and (B) *Hpi*. Trees were inferred from alignments of core genome SNVs, showing relationship between RCH isolates (red tips) and publicly available genome collections (summarised in **Table S1**). Shading indicates strain clusters of RCH isolates involved in potential transmission or persistence (see **Figure 4**).

We used DAPC to explore how the population of RCH CF *Hi* isolates compared to the global *Hi* population structure captured by *k*-mer profiles of publicly available WGS data from a range of other contexts (**File S1, Figure S4**). The data indicate that our Australian paediatric CF isolates are typical of non-invasive respiratory isolates from children in other settings, based on functions constructed to discriminate these parameters from isolates collected from adults, non-respiratory specimens and invasive disease (**Figure 2A-C**). RCH isolates clustered most closely with other Australian isolates in the discriminant function based on geographical location (**Figure 2D**). Notably, DAPC analysis of the 264,940 core-genome SNVs used for phylogenetic inference yielded much weaker discriminant functions for specimen type and geographical location (see **Figure S5**); suggesting the discriminant genetic features are encoded in the accessory genome rather than core gene allelic variation.

**Figure 2.**
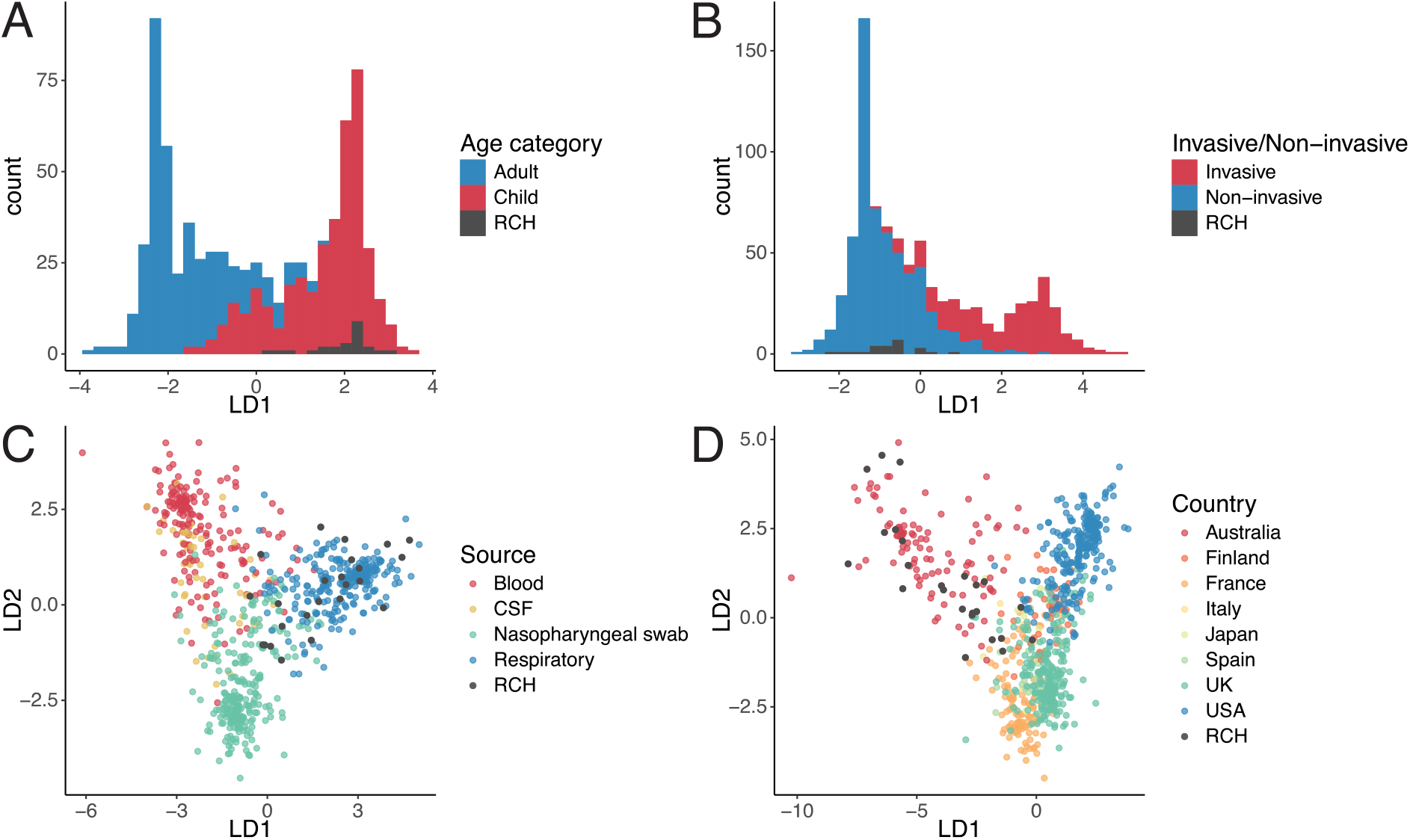
Discriminant analysis of principal components for *Hi* isolates. Analyses were based on *k*-mer profiles extracted from the genomes of novel Australian CF paediatric respiratory colonising isolates sequenced for this study (labelled RCH, black) and those available in public genome collections (other colours; data sources are summarised in **File S1**). Plots show the distribution of values for the significant linear discriminants (LD1, LD2) included in the linear discriminant functions, which were constructed to discriminate genomes on the basis of (A) Host age, (B) Infection status (invasive or non-invasive/colonising), (C) Specimen type, and (D) Country of isolation.

The *Hpi* isolates collected in this study also show extensive genetic diversity (median 5.1% nucleotide divergence in core genome). This is harder to contextualise within the overall species diversity due to the low number of genomes available from other studies but appears to be representative of species-wide phylogenetic diversity (**Figure 1B**).

### Antimicrobial resistance

AMR was relatively common, with 52% of *Hi* and 82% of *Hpi* isolates displaying resistance to at least one of the five drugs tested (**Table 2**). The frequency of co-trimoxazole resistance was similar in both species (35% in *Hi*, 31% in *Hpi*) but ampicillin and/or augmentin resistance was observed at significantly higher rates in *Hpi* compared with *Hi* (*p* < 0.05, see **Table 2**). Rifampicin was observed at low frequencies in both species (17% in *Hi*, 7% in *Hpi*). Multidrug resistance (defined here as resistance to ampicillin or augmentin plus at least one other drug class) was also more commonly detected in *Hpi* although the difference was not statistically significant (see **Table 2**).

**Table 2.** Frequency of non-susceptibility to antimicrobials amongst sequenced isolates. Non-susceptibility was defined as I or R according to clinical breakpoints (see **Methods**). Multidrug resistance (MDR) was defined as resistance to ampicillin or augmentin plus at least one other antimicrobial. Association tests compare resistance rate between species and were performed using Fisher’s exact test (* indicates p < 0.05). Analysis is restricted to isolates that were successfully sequenced and found to be pure cultures, with species identification based on genome data.

|  | *H. influenzae* | *H. parainfluenzae* | Odds ratio (95% CI) | *p*-value |
| --- | --- | --- | --- | --- |
| Ampicillin | 7/23 (30%) | 60/82 (73%) | 6.11 (2.05-20.09) | 0.0004* |
| Augmentin | 1/17 (6%) | 25/79 (32%) | 7.30 (1.02-322.38) | 0.0351* |
| Cefotaxime | 0/23 (0%) | 2/81 (3%) | Inf (0.053-Inf) | 1 |
| Co-trimoxazole | 8/23 (35%) | 25/82 (31%) | 0.82 (0.28-2.55) | 0.8002 |
| Rifampicin | 2/12 (17%) | 4/59 (7%) | 0.37 (0.045-4.61) | 0.2659 |
| Ampicillin or Augmentin | 7/17 (41%) | 62/78 (80%) | 5.41 (1.58-19.72) | 0.0026* |
| One or more drugs tested | 12/23 (52%) | 68/83 (82%) | 4.09 (1.36-12.47) | 0.0058* |
| MDR | 5/23 (22%) | 23/83 (28%) | 1.38 (0.42-5.30) | 0.7897 |

We used the WGS data to explore genetic determinants of AMR in the RCH isolates. Horizontally acquired AMR genes were more frequently found in *Hpi* than *Hi* with 42 (51%) *Hpi* and 5 (21%) *Hi* isolates containing one or more acquired AMR genes (p=0.0107 using Fisher’s exact test; **File S2**). Most common were *bla*_TEM_ genes (36 *bla*_TEM-1_, 4 *bla*_TEM-40_, 2 *bla*_TEM-30_), encoded in the mobile element ICE*Hin* (2 *Hi*, 38 *Hpi*) or small plasmids (3 *Hi*, 2 *Hpi*). Other AMR genes were less common and restricted to *Hpi*, which were generally located in ICE*Hin* elements (11 isolates carried *strAB* [aminoglycosides] and *sul1* [sulfonamides/co-trimoxazole]; 5 *aph3’la* [aminoglycosides]; 1 *tetB* [tetracycline]; and 1 *tetM* [tetracycline] with *msrD* and *mefA* [macrolides]). The resistance cassettes of ICE*Hin* varied in structure and gene content (**Figure 3A**). Four distinct *bla*_TEM_ plasmids were observed and were of similar size (4.3-6.5 kbp). Three were homologous to previously sequenced *Hi* plasmids pLFH64 (2 *Hi*, *bla*_TEM-1_), pA1209 (1 *Hi*, *bla*_TEM-1_), pPN223 (1 *Hpi*, *bla*_TEM-1_); the fourth was a novel plasmid present in one *Hpi* isolate (pM1C124_1, *bla*_TEM-40_) (**Figure 3B** and **Table S3**). Acquired AMR genes only accounted for 33.6% of observed non-susceptibility phenotypes, hence we screened for mutations in core resistance-related chromosomal genes that could potentially explain resistance to ampicillin and augmentin (*pbp* genes, see **Table S2**), co-trimoxazole (*folP, folA*) and rifampicin (*rpoB*) (summarised in **Tables S4–S5**).

**Figure 3.**
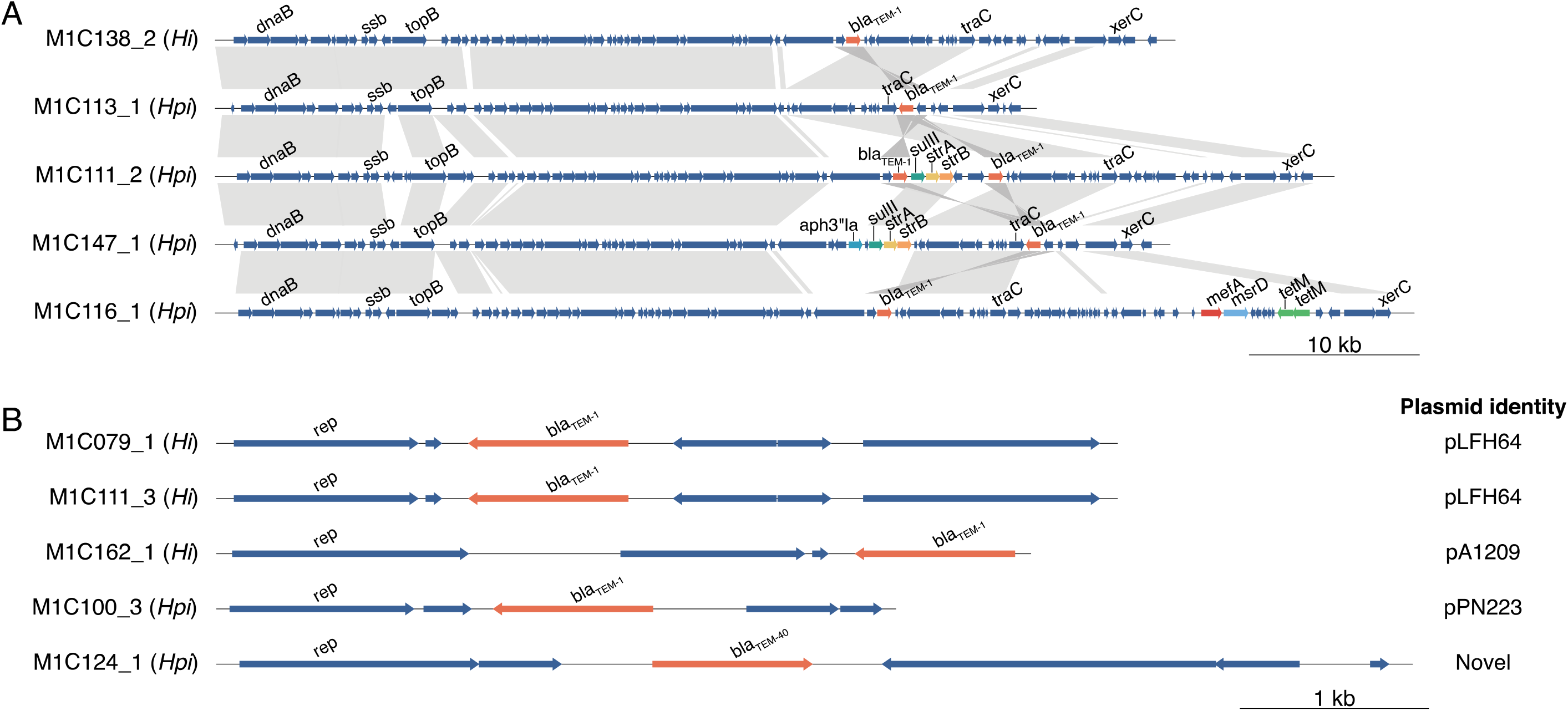
(A) Representative ICE*Hin* structures and (B) plasmids carrying AMR genes identified in RCH isolates. Plasmids are annotated with the best corresponding match in the NCBI nt database, see **Table S3.**

Ampicillin resistance in *Hi* could be entirely explained by acquired β-lactamases encoded by *bla*_TEM_ genes (57%), and/or mutations in the penicillin-binding proteins PBP3 (FtsI) and PBP1B (MrcB) (**Table S4**). In *Hpi*, acquired *bla*_TEM_ could explain 60% of ampicillin resistance, but we detected no known or novel PBP mutation that was statistically associated with resistance (**Table S5**). Notably though, the first *Hpi* isolated from participant M1C152 was ampicillin sensitive and wildtype at PBP3-502, whilst the two subsequent *Hpi* isolates from this participant (following treatment with augmentin and ceftriaxone) were ampicillin resistant with no acquired AMR genes and differed from the first by a single SNV resulting in the amino acid substitution PBP3-A502T, supporting the previously reported role of this mutation in conferring resistance. Nevertheless, this leaves 33% of ampicillin resistance in *Hpi* unexplained.

Augmentin is a combination of amoxycillin plus the β-lactamase inhibitor clavulanic acid. Just four augmentin resistant isolates (16%) carried inhibitor-resistant β-lactamase alleles (2 *Hpi* with *bla*_TEM-30_, 2 *Hpi* with *bla*_TEM-40_); a further nine (36%) carried *bla*_TEM-1_, but this encoded β-lactamase is susceptible to clavulanic acid inhibition and we identified no *pbp* variants in these isolates that were significantly associated with augmentin resistance, hence inhibitor resistance remains unexplained in these cases. Two augmentin resistant *Hpi* isolates collected from the same participant (M1C141) contained novel *bla*_TEM_ alleles that share substitution mutations with known inhibitor-resistant alleles (*bla*_TEM-1_-M67I, *bla*_TEM-1_-W163L) (**Figure S6**), which likely explain the phenotype (34,47). Hence the vast majority of augmentin resistance (100% in *Hi*, 80% in *Hpi*) is unexplained. Resistance to the third-generation cephalosporin cefotaxime was observed in two *Hpi* isolates from different participants but was unexplained (one carried no acquired genes and one carried *bla*_TEM-1_ only, neither carried unique *pbp* mutations).

Co-trimoxazole is a combination of trimethoprim and sulfamethoxazole. Resistance to trimethoprim is associated with mutations in the chromosomal dihydrofolate reductase *folA* or acquisition of mobile resistant alleles (*dfr* genes), while resistance to sulfamethoxazole requires mutations in the chromosomal dihydropteroate synthase *folP* or acquisition of mobile resistant alleles (*sul* genes). In *Hi*, no acquired *sul* or *dfr* genes were detected, however all co-trimoxazole resistant isolates carried a novel resistance-associated mutation FolA-N13S, and most carried the novel FolP-G189C (75%) as well as previously reported FolA-I95L (75%) and FolP-P64ins (38%) (**Table S4, File S2**). In *Hpi*, 92% of co-trimoxazole resistant isolates carried *sul1* (36%) and/or resistance-associated FolP mutations (including FolP-P64ins and FolP-G189C, 80%); 64% carried resistance-associated FolA mutations (38) (**Table S5, File S2**).

Rifampicin resistance is most often explained by mutations in *rpoB* (the RNA polymerase beta subunit), including one previous report in *Hi* (48). Both rifampicin-resistant *Hi* isolates (from the same patient, M1C073) carried a novel mutation RpoB-A1131T that was absent from sensitive isolates. In *Hpi*, one of four rifampicin resistant isolates (M1C081_2) carried RpoB-T724I, which has been previously described in resistant *Hpi* isolates (49), however the remaining three isolates contained no other mutations associated with rifampicin resistance (**File S2**).

### Persistent colonisation and transmission

Seventy-nine participants (53.7%) were culture-positive for the same *Haemophilus* species on ≥1 occasion: 7 (4.8%) had ≥2 *Hi* and 75 (51%) had ≥2 *Hpi*. The probability of testing culture-positive for the same species in the next sample after an initial positive result (mean time interval 105 days) was 16.0% for *Hi* and 48.1% for *Hpi* (*p*-value=0.004 for test of difference in proportions). WGS data was available for 45 such cases and confirmed persistent colonisation with the same strain (1-12 mutations, see **Methods**) in 13 participants (2 *Hi* and 11 *Hpi*, see **Figure 4A**). This represents 8.8% of the total 147 participants, and 29% of the 45 participants that had multiple isolates of the same species available for comparison (95% CI, 16–42%). Assuming this rate of strain matching applied to all 79 participants with ≥2 isolates of the same species, we estimate that 15.6% of the total 147 participants (95% CI, 8.4%–22.6%) had *Haemophilus* colonisation that persisted between visits. Notably the only encapsulated strain detected in this study (cap-e *Hi*) was detected twice in the same participant (M1C094), on two separate clinic visits 84 days apart.

**Figure 4.**
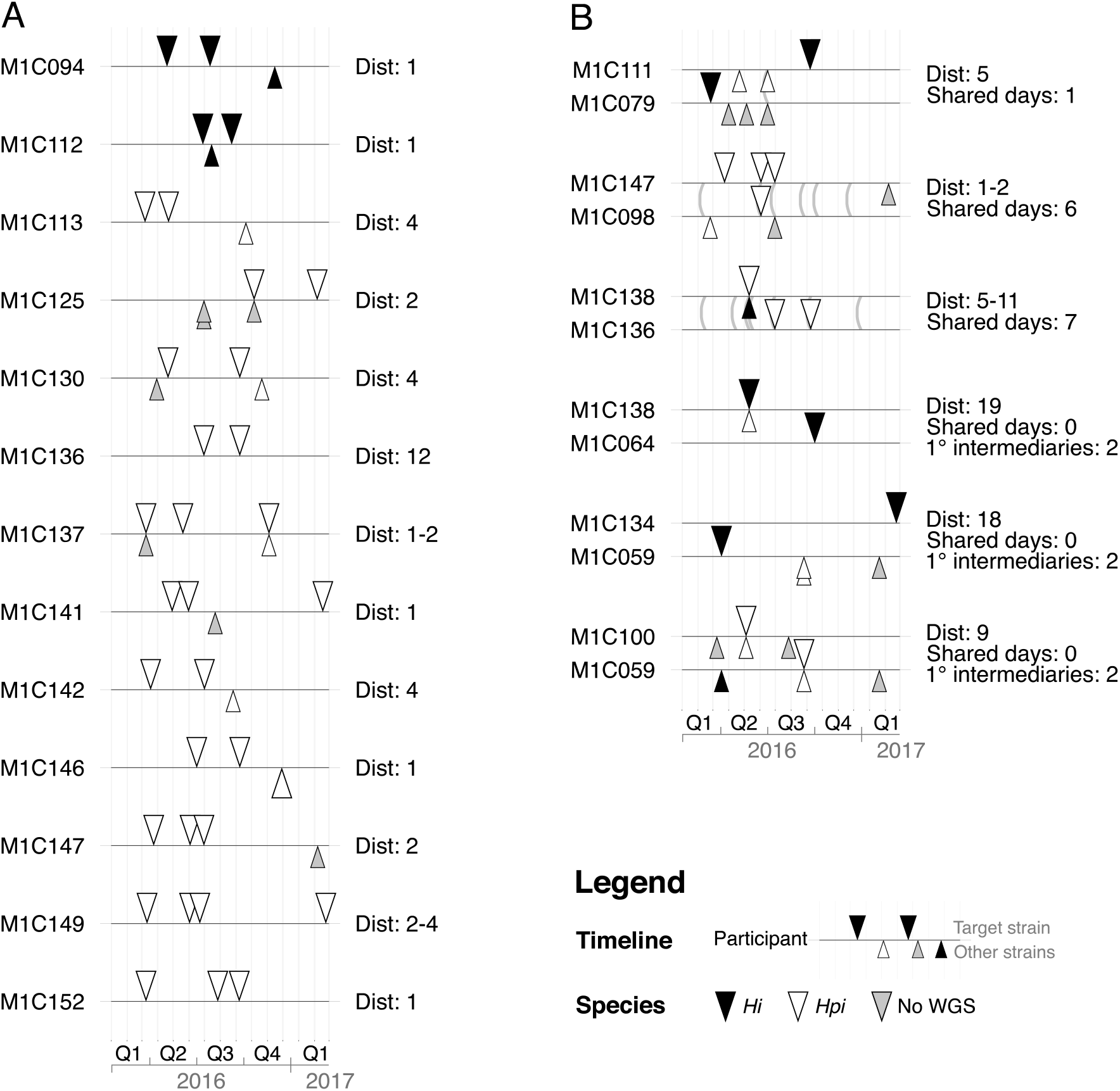
Timelines of *Haemophilus* isolation for participants affected by (A) persistent colonisation or (B) potential transmission between participants. Triangles indicate *Haemophilus* isolates, coloured according to WGS-confirmed species as per inset legend. Isolates presented above each timeline represent the same strain (defined as pairwise genetic distance <20, see **Methods**), those below are different strains. Dist: range of pairwise genetic distances (non-recombinant SNVs + number of inferred recombination events) observed between isolates of the same strain. In (B), lines connecting participant timelines represent instances where participants attended an RCH clinic during the same day. Shared days: number of days on which both participants attended the RCH CF clinic; 1° intermediaries: for participant pairs not sharing any clinic visit days with one another, we searched for primary intermediary participants who shared at least one clinic data with each of the participants.

WGS data showed most strains were unique to a single participant, however we identified six participant pairs that shared the same strain (1-19 mutations, see **Methods**; 3 *Hi*, 3 *Hpi*, see **Figure 4B**), suggesting potential transmission. In three such cases, both participants visited an RCH CF clinic on the same day (**Figure 4B**), providing a potential opportunity for transmission within the clinic. A contact network reconstructed from visit dates showed that while the other three strain-sharing participant pairs never attended on the same day, they did have shared visit days with single-step intermediary participants (**Figure S7**); hence it is possible that one of these intermediary participants had become colonised at the clinic via the first participant, and then passed the strain on to the second participant on a subsequent visit.

Twelve of the thirteen (92%) WGS-confirmed cases of persistent strain colonisation involved resistance to at least one drug, compared with 70% of strains that were not identified as persisting (see **Table S6, File S2**). For *Hpi*, all AMR phenotypes were more frequent amongst isolates associated with persistent strain colonisation, however these comparisons were underpowered and the differences were only statistically significant for ampicillin, augmentin, and cefotaxime (**Table S6**). Five of the six potentially transmitted strains displayed at least one AMR phenotype, similar to the overall rate of AMR across all colonising isolates (**Table S7**, **File S2**). Changes in AMR phenotypes within individual *Hi* or *Hpi* strains were observed during both persistent colonisation (resistance phenotypes varied in 8/13 individuals, 62%) and transmission chains (resistance phenotypes varied in 4/6 transmission pairs, 67%), indicating short-term evolution of resistance (**File S2**).

## Discussion

*Hi* and *Hpi* colonisation of the airways was strikingly common in this cohort, with >80% of participants contributing ≥1 *Haemophilus* culture-positive respiratory sample during the 1-year period of study (**Table 1**) and 58.5% contributing ≥2 such samples. Point prevalence was 4.6% for *Hi* and 32.1% for *Hpi*, and repeat colonisation with *Hpi* was much more common than *Hi* (detected in 51% and 4.8% of participants, respectively). *Hpi* also displayed a significantly higher frequency of AMR (**Table 2**), perhaps linked to its increased carriage rate and the associated increase in exposure to antimicrobials. Globally the *Hi* carriage rate in childhood cohorts varies, likely due to differences in cohort demographics and geographical location. *Hi* has been reported to be recoverable from the nasopharynx in 8% to 34% of children with CF (50–54,20), consistent with *Hi* carriage estimation in this CF cohort. The frequency of *Hpi* airway colonisation has not been established in children (with or without CF) despite the potential to opportunistically cause disease and act as a reservoir for AMR genes.

Substantial genetic diversity was observed for both *Hi* and *Hpi* isolates cultured from the airways of participants in this study (**Figure 1**). Unsurprisingly, only two *Hi* isolates (from a single patient) were predicted to be encapsulated (*cap-e*); these belonged to a known clonal capsule-positive lineage (ST18). The remaining *Hi* isolates belong to the highly heterogenous NTHi group, similar to those detected in other studies examining nasopharyngeal colonisation, which consistently report NTHi as the dominant *Hi* subtype in the respiratory tract (20,21).

Analysis of *Hi* core-genome SNVs using phylogenetics and DAPC showed no apparent lineage associations with age group, specimen type, disease status or geographical location (**Figure S4** and **S5**), while whole-genome *k*-mer DAPC revealed distinct clustering of RCH CF isolates with others that were non-invasive, collected the respiratory tract, isolated from children, and circulating in Australia (based on the respective individual discriminant functions, see **Figure 2**). Hence *Hi* isolates of distinct epidemiological origins are differentiable based on variation in accessory genes, but not by allelic variation in the core genome.

The structured variability across the accessory genome could potentially be explained by niche-specific positive selection of genes that confer increased fitness. For example, fixation of ICE*Hin*1056 in respiratory *Hi* populations has been previously observed within two weeks of amoxicillin treatment, but subsequently lost (or resistant strains outcompeted) twelve weeks after the initial treatment (55). The Hia and HMW adhesins, Hif pilus, and IgA-proteases are *Hi* virulence factors that are also differentially present in strains (56,57) and play a role in colonisation of specific niches like the respiratory tract (58–61). Genome-wide association analysis could potentially identify other contributing factors (62), however this is beyond the scope of the present study.

Antimicrobial therapy is used routinely both to control bacterial lung infections of CF patients (63) and also as antimicrobial prophylaxis. Augmentin is routinely used for both these purposes. Co-trimoxazole is used at many specialist CF centres but it is not routine at RCH, however resistance was still observed in nearly a third of *Hi* and *Hpi*. Regular use of antimicrobials is known to induce resistance and indeed we observed a high rate of AMR in isolates collected in this study, with resistance to one or more drugs observed in 52% *Hi* and 82% *Hpi* (**Table 2**). AMR rates in *Hi* isolated from the respiratory tract varies between studies, with recent reports of ampicillin resistance 23.7% to 58.5%, augmentin 0% to 10.4%, cefotaxime 0% to 5.9%, co-trimoxazole 21.4% to 71.1%, and rifampicin 0% and 4.8% (49,64–72). Similar rates have been reported for *Hpi:* ampicillin 13.2% to 50.0%, augmentin 0% to 12.5%, cefotaxime 0% to 0.3%, co-trimoxazole 14.9% to 44.2%, and rifampicin 1.5% to 26.7% (34,73–76). AMR rates detected in the present study are mostly in line with these reports, with the exception of higher rates of resistance in *Hpi* vs *Hi*. The only other report of such a difference is an earlier study in our setting (children with CF at RCH, 1998-2012) (75), which also found higher rates of resistance in *Hpi* vs *Hi*, and showed that rates of ampicillin, augmentin and co-trimoxazole resistance increased significantly in *Hpi* over the 15-year duration of the study.

Most of the AMR phenotypes were explained by the presence of known genetic determinants. Ampicillin resistance was the most readily explained by known mechanisms, with all *Hi* resistant isolates and 67% of *Hpi* resistant isolates harbouring the acquired gene *bla*_TEM_ or resistance-associated mutations in PBP genes (**File S2**). The exceptions were augmentin and ceftriaxone: inhibitor-resistant *bla*_TEM_ alleles accounted for just 20% of augmentin resistance in *Hpi* and none in *Hi*, and no mechanisms for ceftriaxone resistance were identified.

Novel mutations in AMR-associated proteins discovered through association analysis increased the proportion of resistance explained by amino acid substitutions from 13.4% to 31.3%. Several mutations in FolA and FolP were associated with co-trimoxazole resistance (**Table S4-S5**), including both novel and previously established mutations (38). Notably we identified an insertion in *Hpi* FolP that was strongly linked with co-trimoxazole resistance and shared the same location as the *Hi* FolP-P64ins mutation, which has been demonstrated to induce sulfamethoxazole resistance (33). Mutations for rifampicin resistance were identified in *Hi* (RpoB-A1131T) and *Hpi* (RpoB-T724I); the latter was also recently reported as resistance-associated in an independent study of *Hpi* (49).

Consistent with previous studies, nearly all acquired genes detected here in *Haemophilus* isolates were localised to either a ICE*Hin* element (77,78) or small *bla*_TEM_ plasmids (14,34,79). Novel variants of acquired resistant determinants were also observed, including a new ICE*Hin*-encoded *bla*_TEM_ allele associated with augmentin resistance in *Hpi*, and a novel plasmid harbouring *bla*_TEM-1_.

Not all AMR could be explained by an underlying genetic component. This is likely due in part to a lack of statistical power to detect novel resistance-associated variants, even when taking a candidate-gene approach as we did, due to small sample size. This is particularly problematic for *Hi*, for which only 24 sequenced isolates were available; for example, FolP-G189C was associated with co-trimoxazole resistance in both *Hi* and *Hpi* but was statistically significant only in *Hpi* after adjustment for multiple testing (**Table S4-S5**). Additionally, AMR phenotypes are not always reproducible and AMR genes or mutations can be lost during subculture to extract DNA for sequencing.

There is little published data on persistence of *Haemophilus* colonisation in the lungs of children, however there is some evidence that *Hi* strain persistence is associated with chronic respiratory disease and does not occur in healthy childhood cohorts (80). This study reveals for the first time strain persistence of *Hi* and *Hpi* in the lungs of children with cystic fibrosis for up to 349 days, and estimates carriage rate of persistent strains in the cohort to be 15.6% (95% CI, 8.4%-22.6%). Strain persistence likely arises due to substantial selective pressure exerted by extensive and prolonged administration of antimicrobials or niche adaptation to the diseased lung. For example, mutations in the single-strand mispairing mechanism allow *Hi* to alter expression of nutrient uptake systems and surface molecules such as adhesins during persistent colonisation in adult patients with chronic obstructive pulmonary disorder (81). Other important CF pathogens such as *P. aeruginosa* and members of the *B. cepacia* complex have been shown to undergo similar changes to surface molecules and remodelling of regulatory networks during persistence (82–84). In addition to strain persistence, we observed that participants were frequently colonised by different strains of *Hi* and *Hpi* across clinic visits, indicating that *Haemophilus* colonisation is a dynamic process and suggesting that strains of both species compete to occupy the niche.

There were six instances where participants shared the same *Haemophilus* strain and three of the six cases were supported by epidemiological links whereby participants shared clinic visit days (**Figure S7**), and the remaining three cases shared visit days with possible intermediaries (**Figure S7**). Nosocomial transmission of CF pathogens such as *P. aeruginosa* has been demonstrated in other settings (84,85) and historically at our centre (86,87), as has cross-infection with *Mycobacterium abscessus* (88). This has led to strict infection control practices in CF clinics such as RCH with strict isolation in both clinics and in-patient areas, wearing of face-masks by patients in all public spaces and strict gloving and gowning for by all clinical staff. Notably, preceding the introduction of such stringent infection-control measures, sharing of RCH CF clinic visit days was common in our cohort and nearly all participants pairs could be connected either through a shared clinic visit day (15%) or shared visit days with a single intermediary (72.4%), hence it is not clear whether the overlap in visit days could be circumstantial and strain sharing may reflect circulation of strains in the general community rather than nosocomial transmission.

This study provides the first insights into the population dynamics and genomic determinants of AMR amongst colonising *Hi* and *Hpi* strains in a paediatric CF cohort, and identifies multiple novel AMR determinants particularly for *Hpi*. Notably, while relatively little attention has been paid to *Hpi* colonisation in children due to its relative lack of pathogenicity, our data indicate it is a common coloniser that can persist in the respiratory tract of CF children and is very frequently drug resistant. The high frequency of AMR in *Hpi*, most of which was encoded in mobile elements that can transfer to *Hi*, indicates that *Hpi* could serve as a reservoir for the emergence and spread of AMR to *Hi* which is of more significant clinical concern in children with and without CF. Further insights are essential and will inform antimicrobial treatment and stewardship in the future.

## Acknowledgments

This work was supported by the Bill & Melinda Gates Foundation, Seattle (KEH) and the Australian Government Research Training Program (SW). KEH is supported by a Senior Medical Research Fellowship from the Viertel Foundation of Victoria. All the authors declare that there are no conflicts of interest.

## Supporting Files

**File S1**. Isolates sequenced in this study.

**File S2**. AMR phenotypes of isolates and genetic mechanisms of resistance.

## Supporting Figures

**Figure S1.**
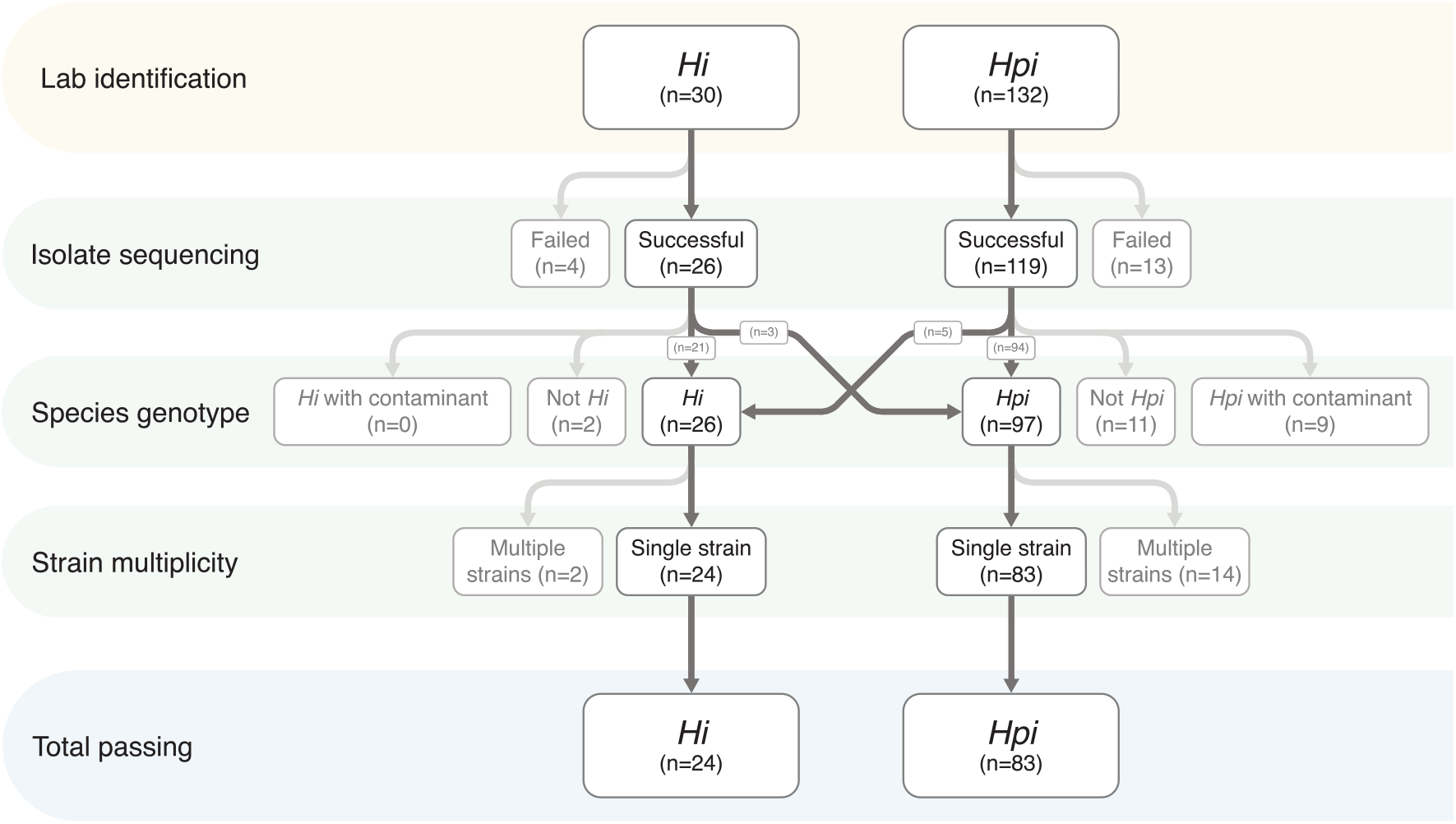
Quality control applied to read data and isolates passing each stage.

**Figure S2.**
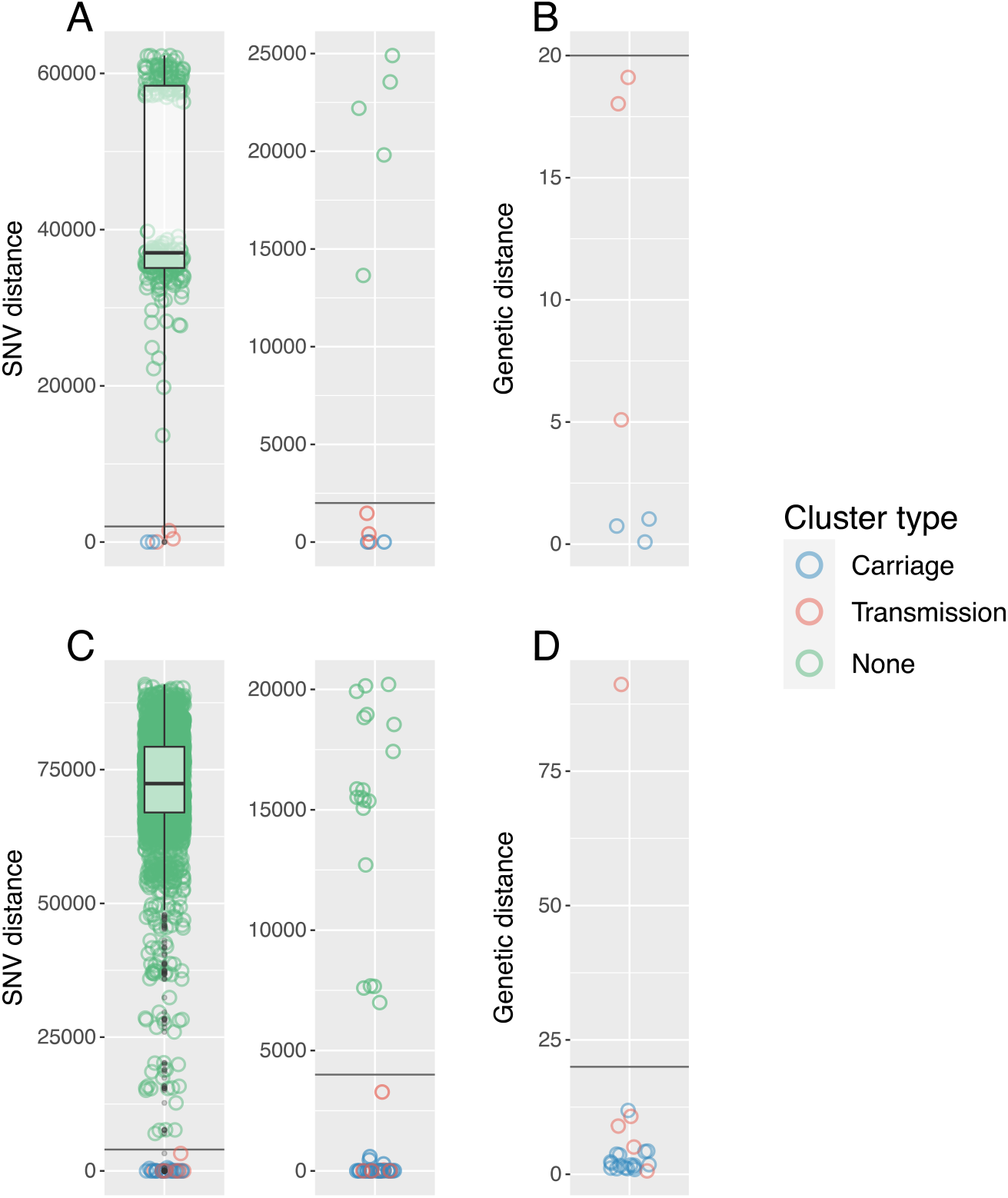
Distributions of pair-wise distances between isolates sequenced in this study. Core genome SNV distances shown for all isolate pairs and isolate pairs ≤25,000 SNVs for (A) *Hi* and (C) *Hpi*. Isolates passing the species-specific SNV distance threshold (*Hi*: 2,000; *Hpi*: 4,000) were mapped to their respective cluster reference and a new, more accurate genetic measure was defined as non-recombinant SNVs + number of recombination events (see **Methods**) for (B) *Hi* and (D) *Hpi*. Isolate pairs with a genetic distance <20 were considered the same strain. Black lines indicate the applied thresholds.

**Figure S3.**
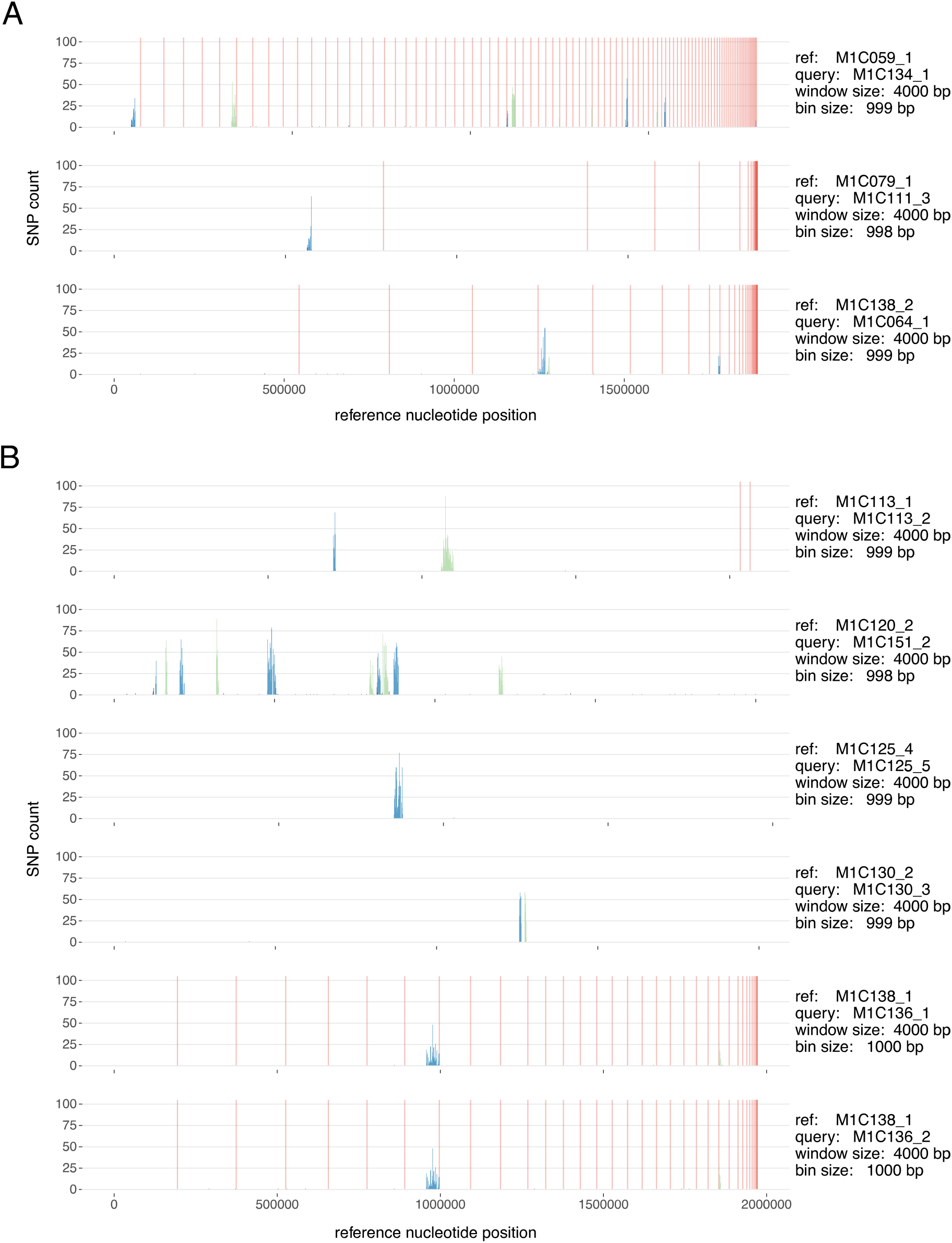
Histograms of core SVNs in transmission and carriage isolates with detected recombination for (A) *H. influenzae* and (B) *H. parainfluenzae* (SVNs are relative to the nominated reference). Blocks of SVNs significantly denser than background (and thus a product of recombination) are coloured alternating blue and green (*p*-value ≤ 0.05, Bonferroni-corrected). Black histogram bars represent non-recombinant SVNs and vertical red lines denote contig boundaries in the nominated reference.

**Figure S4.**
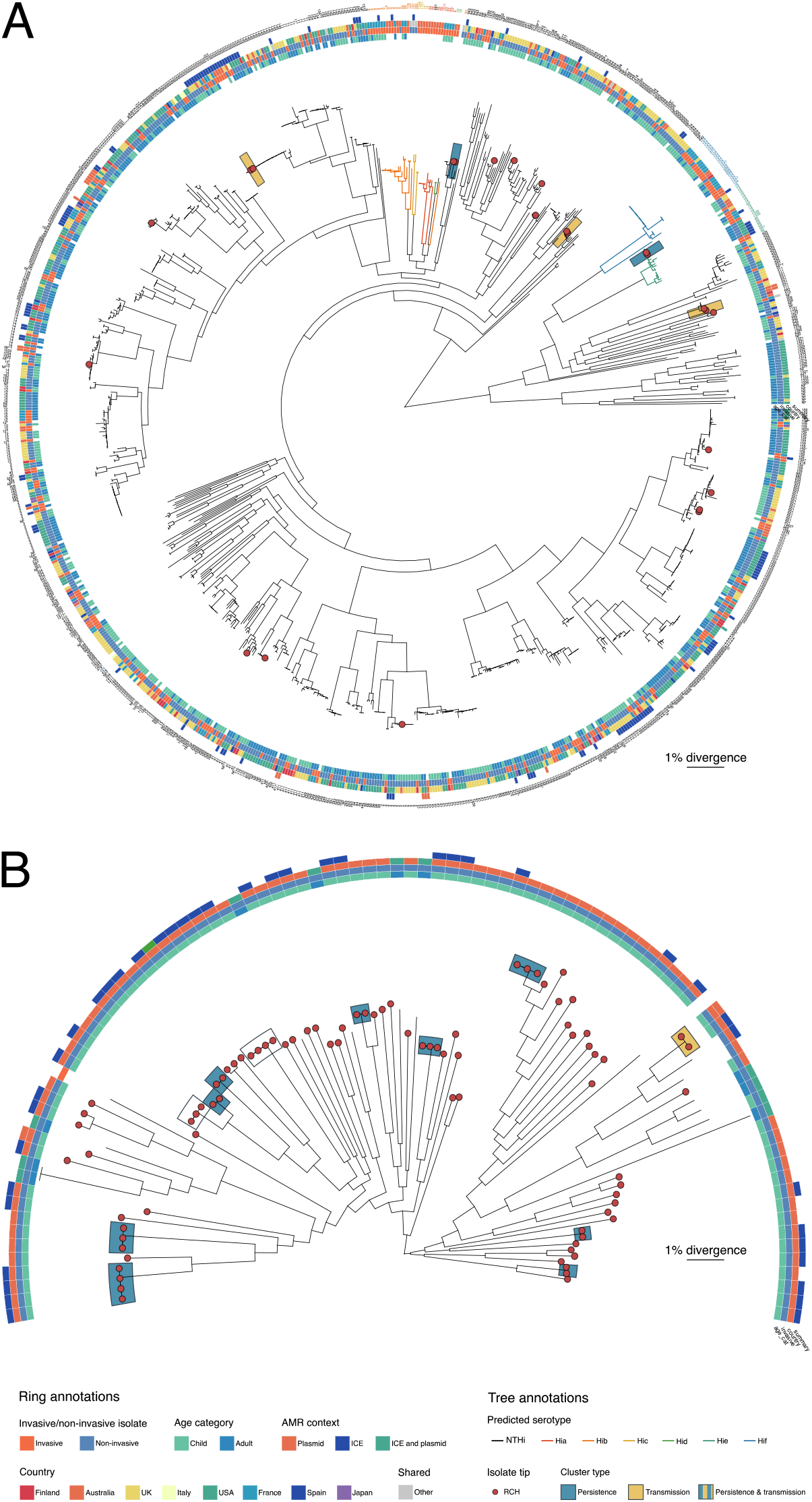
Maximum-likelihood phylogenies annotated with sample information for (A) *Hi* and (B) *Hpi*. Trees were inferred from alignments of core genome SNVs, showing relationship between RCH isolates (red tips) and publicly available genome collections (summarised in **Table S1**). Shading indicates strain clusters of RCH isolates involved in potential transmission or persistence (see **Figure 4**).

**Figure S5.**
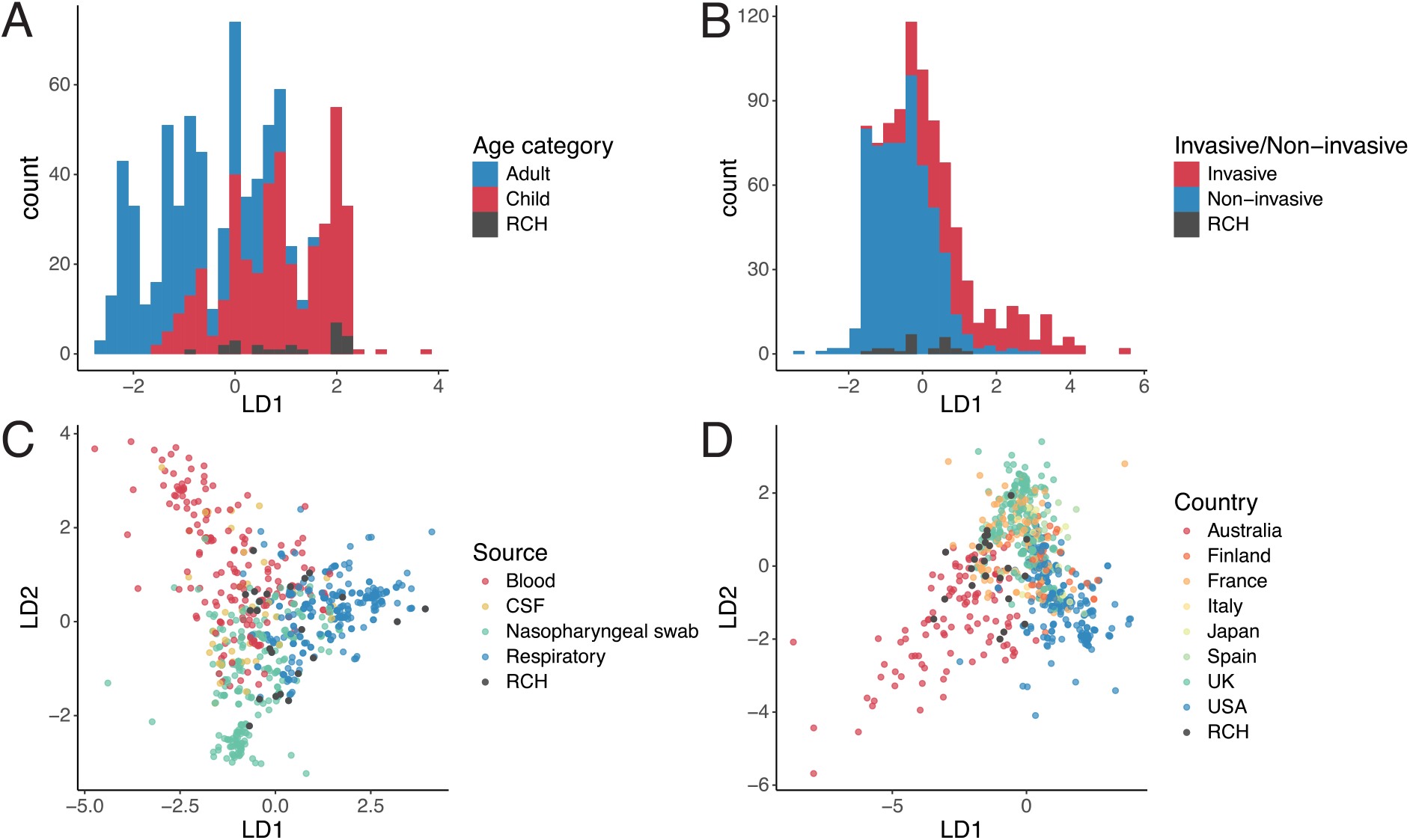
Discriminant analysis of principal components for *Hi* isolates. Analyses were based on core SNVs from the genomes of novel Australian CF paediatric respiratory colonising isolates sequenced for this study (labelled RCH, black) and those available in public genome collections (other colours; data sources are summarised in **File S1**). Plots show the distribution of values for the significant linear discriminants (LD1, LD2) included in the linear discriminant functions, which were constructed to discriminate genomes on the basis of (A) Host age, (B) Infection status (invasive or non-invasive/colonising), (C) Specimen type, and (D) Country of isolation.

**Figure S6.**
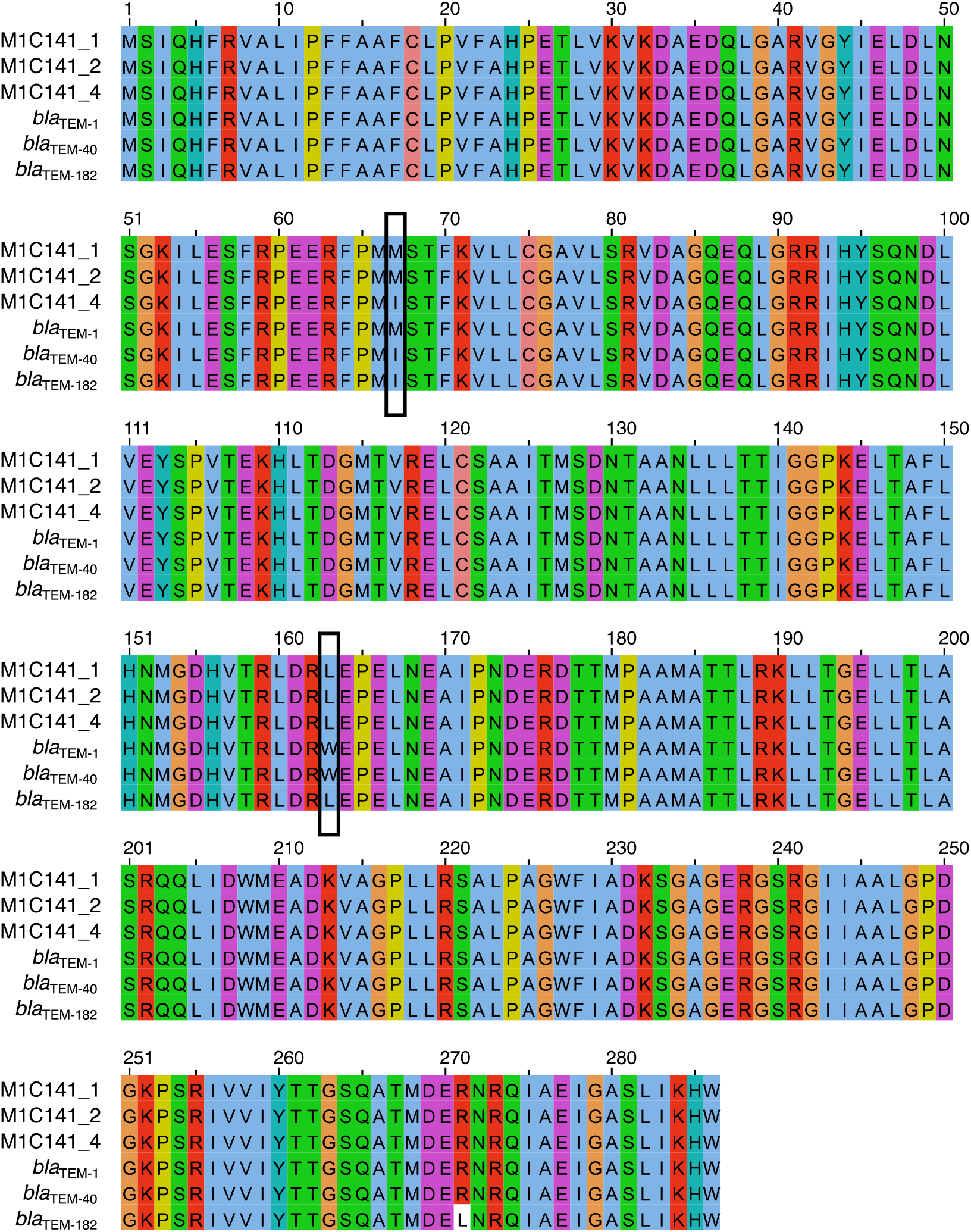
Multiple sequence alignment of *bla*_TEM_ alleles from M1C141 isolates and reference *bla*_TEM-1_, *bla*_TEM-40_, and *bla*_TEM-182_ from the NCBI AMR database. The *bla*_TEM-40_ and *bla*_TEM-182_ alleles are inhibitor resistance, and *bla*_TEM-1_ is inhibitor sensitive. Variants in M1C141 *bla*_TEM_ alleles relative to reference sequences are indicated by black boxes.

**Figure S7.**
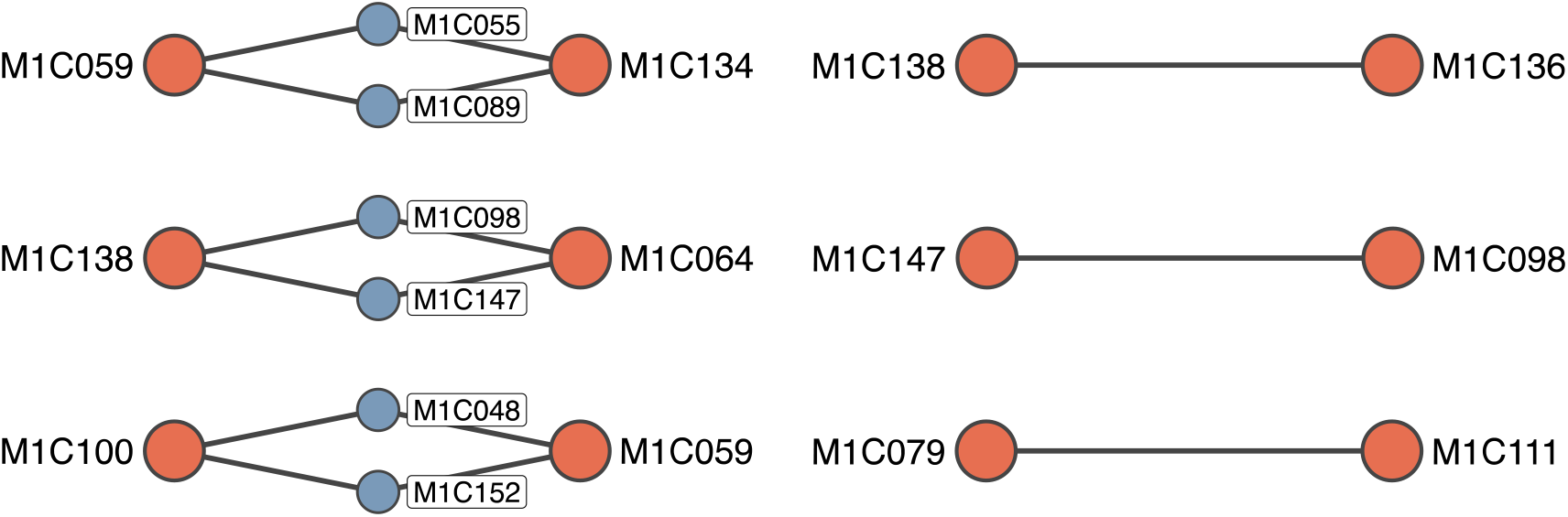
Shortest traversal paths between transmission host endpoints extracted from hospital visitation contact network. Nodes are participants and edges indicate participants that shared at least one visitation day after initial detection of first transmission isolate.

## Supporting Tables

**Table S1.** Public datasets used in this study.

| Species | Study | Genomes | DOI | Data source |
| --- | --- | --- | --- | --- |
| *Hi* | Price *et al.* 2015 | 2 | https://doi.org/10.1186/s12864-015-1857-x | PRJNA292146 |
|  | Giufre' *et al.* 2015 | 11 | https://doi.org/10.1128/genomeA.00110-15 | PRJNA272685 |
|  | Naito *et al.* 2018 | 14 | http://dx.crossref.org/10.1128/JCM.00141-18 | PRJDB7020 |
|  | Staples *et al.* 2017 | 98 | https://doi.org/10.1017/S0950268817000450 | PRJEB18702 |
|  | De Chiara *et al.* 2014 | 89 | https://doi.org/10.1073/pnas.1403353111 | Sanger FTP<sup>a</sup> |
|  | Cleary *et al.* 2018 | 265 | https://dx.doi.org/10.1099%2Fmgen.0.000209 | PRJEB23674 |
|  | Pettigrew *et al.* 2018 | 270 | https://doi.org/10.1073/pnas.1719654115 | PRJNA358390 |
|  | Deghmane *et al.* 2019 | 128 | https://doi.org/10.1016/j.jinf.2019.05.007 | PRJEB28646 |
| *Hpi* | Price *et al.* 2017 | 3 | https://doi.org/10.2217/fmb-2016-0215 | PRJNA327273 |
|  | Roach *et al.* 2015 | 11 | https://dx.doi.org/10.1371%2Fjournal.pgen.1005413 | PRJNA267549 |

<sup>a</sup> ftp://ftp.sanger.ac.uk/pub/project/pathogens/Haemophilus/influenzae/NT_strains/

**Table S2.** PBP homologs identified in reference genomes *Hi* strain Rd KW20 and *Hpi* strain T3T1. Reference proteins were aligned to curated PBP protein sequences in the Swiss-Prot database. Proteins were considered homologs where the respective alignment had ≥70% identity and ≥80% coverage.

| Protein | Reference genome gene identifier |  |
| --- | --- | --- |
|  | *H. influenzae* | *H. parainfluenzae* |
| PBP1A | HI0440 | PARA_RS04050 |
| PBP1B (MrcB) | HI1725 | PARA_RS08825 |
| PBP2 (MrdA) | HI0032 | PARA_RS00610 |
| PBP3 (FtsI) | HI1132 | PARA_RS06305 |
| PBP4 (DacB) | HI1330 | PARA_RS08695 |
| PBP5 (DacA) | HI0029 | PARA_RS00625 |
| PBP7 | HI0364 | PARA_RS00190 |
| MepA | HI0197 | PARA_RS09660 |

**Table S3.** Best alignment matches of isolate plasmids found in the NCBI nt database using BLASTn. The plasmid from M1C124_1 had no close match in the database (low coverage indicated in bold typeface).

| Isolate | Species | Plasmid name | Identity (%) | Coverage (%) |
| --- | --- | --- | --- | --- |
| M1C079_1 | *Hi* | pLFH64 | 99.75 | 100.00 |
| M1C111_3 | *Hi* | pLFH64 | 99.75 | 100.00 |
| M1C162_1 | *Hi* | pA1209 | 100.00 | 100.00 |
| M1C100_3 | *Hpi* | pPN223 | 93.73 | 100.00 |
| M1C124_1 | *Hpi* | pJ612 | 99.93 | **23.15** |

**Table S4.**
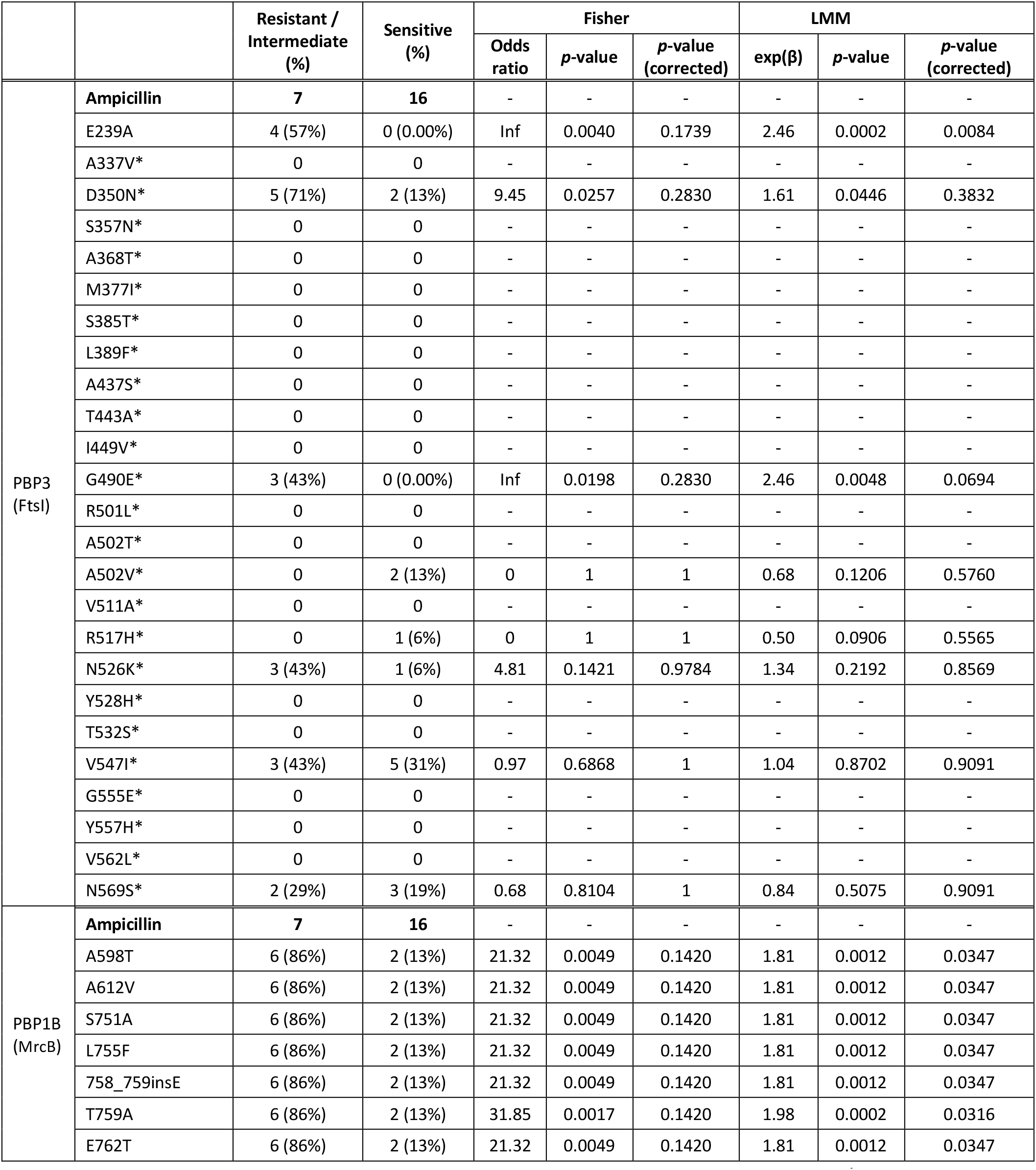
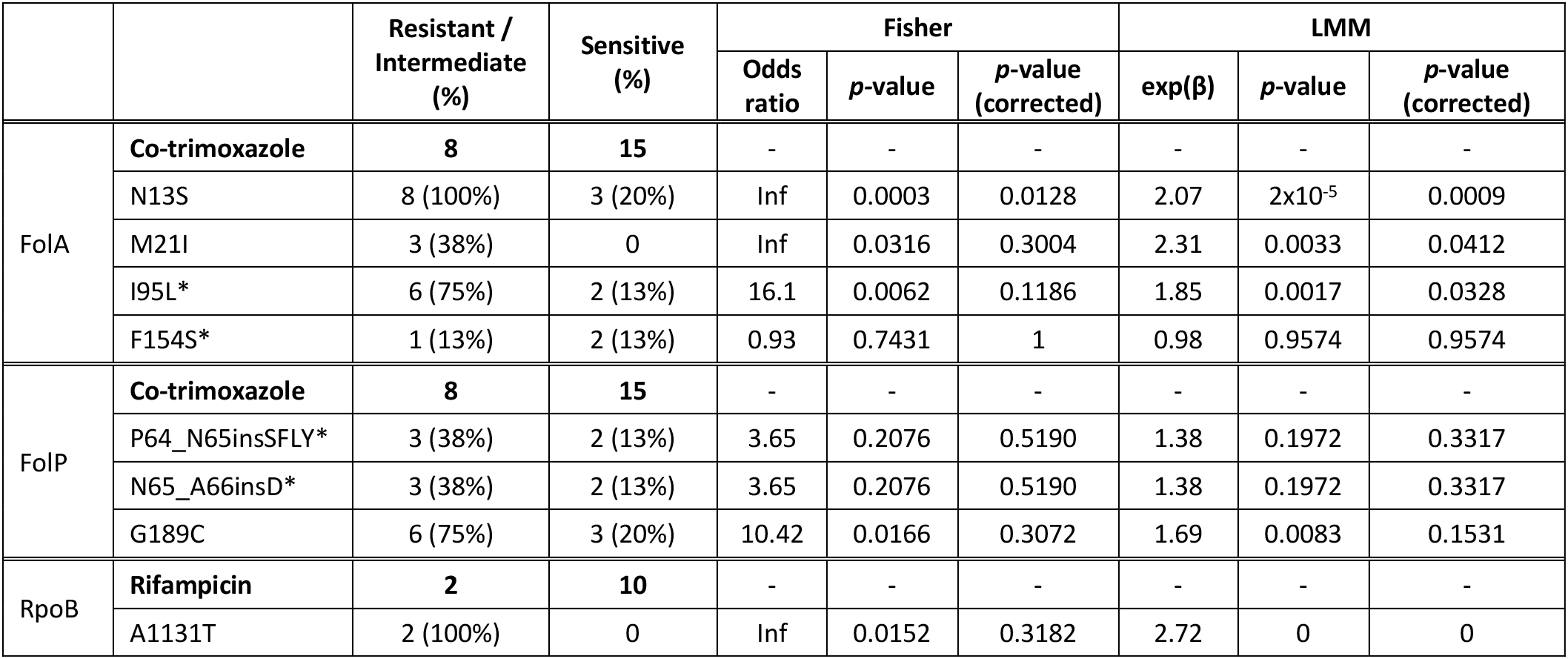
Mutations detected/investigated in AMR-associated proteins encoded in the *H. influenzae* genome. All mutations previously reported to be associated with the phenotype are included and indicated with *. Novel mutations that were not significantly associated with the respective phenotype (*p*-value ≥ 0.05 in all tests) are not shown. See **Methods** and **Table S2** for full list of proteins investigated.

**Table S5.** Mutations detected/investigated in AMR-associated proteins encoded in the *H. parainfluenzae* genome. All mutations previously reported to be associated with the phenotype are included and indicated with *. Novel mutations that were not significantly associated with the respective phenotype (*p*-value ≥ 0.05 in all tests) are not shown. See **Methods** and **Table S2** for full list of proteins investigated.

|  |  | Resistant / Intermediate (%) | Sensitive (%) | Fisher |  |  | LMM |  |  |
| --- | --- | --- | --- | --- | --- | --- | --- | --- | --- |
| | | | | Odds ratio | $p$-value | $p$-value (corrected) | exp($\beta$) | $p$-value | Odds ratio |
| PBP3<br>(ftsI) | **Ampicillin** | **60** | **22** | - | - | - | - | - | - |
|  | A337V* | 0 | 0 | - | - | - | - | - | - |
|  | S357N* | 1 (2%) | 0 | Inf | 0.7284 | 1 | 1.3 | 0.5590 | 0.8054 |
|  | M377I* | 0 | 0 | - | - | - | - | - | - |
|  | S385T* | 1 (2%) | 1 (5%) | 0.37 | 0.9287 | 1 | 0.78 | 0.4343 | 0.8054 |
|  | L389F* | 0 | 0 | - | - | - | - | - | - |
|  | A437S* | 0 | 0 | - | - | - | - | - | - |
|  | T443A* | 0 | 0 | - | - | - | - | - | - |
|  | I449V* | 0 | 0 | - | - | - | - | - | - |
|  | G490E* | 0 | 0 | - | - | - | - | - | - |
|  | R501L* | 0 | 0 | - | - | - | - | - | - |
|  | A502V* | 3 (5%) | 1 (5%) | 1.12 | 0.7034 | 1 | 1.01 | 0.9817 | 0.9939 |
|  | A502T* | 3 (5%) | 0 | Inf | 0.3810 | 1 | 1.39 | 0.2217 | 0.8054 |
|  | R517H* | 1 (2%) | 0 | Inf | 0.7284 | 1 | 1.3 | 0.5590 | 0.8054 |
|  | Y528H* | 0 | 0 | - | - | - | - | - | - |
|  | T532S* | 0 | 0 | - | - | - | - | - | - |
|  | G555E* | 0 | 0 | - | - | - | - | - | - |
|  | Y557H* | 1 (2%) | 0 | Inf | 0.7284 | 1 | 1.3 | 0.5590 | 0.8054 |
|  | V562L* | 2 (3%) | 1 (5%) | 0.74 | 0.8222 | 1 | 0.93 | 0.7927 | 0.9539 |
| FolA | **Co-trimoxazole** | **25** | **57** | - | - | - | - | - | - |
| | W31R | 10 (40%) | 2 (4%) | 17.51 | $7 \times 10^{-5}$ | 0.0022 | 1.84 | $8 \times 10^{-5}$ | 0.0018 |
| | I95L* | 13 (52%) | 2 (4%) | 28.09 | $9 \times 10^{-7}$ | $6 \times 10^{-5}$ | 2.07 | $2 \times 10^{-7}$ | $1 \times 10^{-5}$ |
| | F154V* | 6 (24%) | 1 (2%) | 16.99 | 0.0028 | 0.0612 | 2.12 | $4 \times 10^{-5}$ | 0.0014 |
|  | F154S* | 5 (20%) | 2 (4%) | 6.69 | 0.0251 | 0.1840 | 1.56 | 0.0174 | 0.1461 |
| FolP | **Co-trimoxazole** | **25** | **57** | - | - | - | - | - | - |
| | F14L | 15 (60%) | 4 (7%) | 18.12 | $1 \times 10^{-6}$ | $2 \times 10^{-5}$ | 1.7 | $6 \times 10^{-7}$ | $9 \times 10^{-6}$ |
| | S31G | 20 (80%) | 5 (9%) | 36.78 | $5 \times 10^{-10}$ | $2 \times 10^{-8}$ | 1.96 | $3 \times 10^{-11}$ | $1 \times 10^{-9}$ |
| | P64_M65insI | 20 (80%) | 5 (9%) | 36.78 | $5 \times 10^{-10}$ | $2 \times 10^{-8}$ | 1.96 | $3 \times 10^{-11}$ | $1 \times 10^{-9}$ |
| | D167A | 20 (80%) | 16 (28%) | 9.43 | $2 \times 10^{-5}$ | 0.0003 | 1.43 | 0.0007 | 0.009 |
| | Q170K | 19 (76%) | 5 (9%) | 29.42 | $4 \times 10^{-9}$ | $9 \times 10^{-8}$ | 1.89 | $2 \times 10^{-9}$ | $5 \times 10^{-8}$ |
| | A174D | 19 (76%) | 6 (11%) | 24.2 | $1 \times 10^{-8}$ | $3 \times 10^{-7}$ | 1.84 | $3 \times 10^{-8}$ | $5 \times 10^{-7}$ |
| | G189C | 20 (80%) | 5 (9%) | 36.78 | $5 \times 10^{-10}$ | $2 \times 10^{-8}$ | 1.96 | $3 \times 10^{-11}$ | $1 \times 10^{-9}$ |
| RpoB | **Rifampicin** | **4** | **55** | - | - | - | - | - | - |
| | T724I | 1 (25.00%) | 0 | Inf | 0.0678 | 1 | 2.58 | $1 \times 10^{-4}$ | 0.0031 |

**Table S6.** AMR frequencies in strains responsible for persistent colonisation compared to singleton strains. Nonsusceptibility was defined as I or R according to clinical breakpoints (see **Methods**). MDR was defined as nonsusceptibility to ampicillin or augmentin plus at least one other antimicrobial. Associations were calculated using two-sided Fisher’s exact test. Significant associations (p < 0.05) are indicated by bold typeface.

|  | *H. influenzae* |  |  |  | *H. parainfluenzae* |  |  |  | Both |  |  |  |
| --- | --- | --- | --- | --- | --- | --- | --- | --- | --- | --- | --- | --- |
|  | Persistent colonisation | Other | OR (95%CI) | *p*-value | Persistent colonisation | Other | OR (95%CI) | *p*-value | Persistent colonisation | Other | OR (95%CI) | *p*-value |
| Ampicillin | 0/2<br>(0%) | 7/19<br>(37%) | 0.00<br>(0.00-10.88) | 0.5333 | 11/11<br>(100%) | 37/55<br>(67%) | Inf<br>(1.06-Inf) | **0.0276** | 11/13<br>(85%) | 44/74<br>(59%) | 3.70<br>(0.73-36.72) | 0.1201 |
| Augmentin | 0/2<br>(0%) | 1/13<br>(8%) | 0.00<br>(0.00-252.50) | 1 | 7/11<br>(64%) | 14/52<br>(27%) | 4.62<br>(1.00-25.01) | **0.0323** | 7/13<br>(54%) | 15/65<br>(23%) | 3.81<br>(0.94-16.13) | **0.0401** |
| Cefotaxime | 0/2<br>(0%) | 0/19<br>(0%) | 0.00<br>(0.00-Inf) | 1 | 2/11<br>(18%) | 0/53<br>(0%) | Inf<br>(0.95-Inf) | **0.0273** | 2/13<br>(15%) | 0/72<br>(0%) | Inf<br>(1.08-Inf) | **0.0218** |
| Co-trimoxazole | 1/2<br>(50%) | 6/19<br>(32%) | 2.08<br>(0.024-182.62) | 1 | 5/11<br>(45%) | 14/54<br>(26%) | 2.35<br>(0.49-10.95) | 0.2753 | 6/13<br>(46%) | 20/73<br>(27%) | 2.25<br>(0.55-8.93) | 0.1995 |
| Rifampicin | 0/2<br>(0%) | 2/8<br>(25%) | 0.00<br>(0.00-26.01) | 1 | 2/10<br>(20%) | 2/41<br>(5%) | 4.67<br>(0.30-73.67) | 0.1682 | 2/12<br>(17%) | 4/49<br>(8%) | 2.21<br>(0.18-18.13) | 0.5879 |
| Ampicillin or Augmentin | 0/2<br>(0%) | 7/13<br>(54%) | 0.00<br>(0.00-6.00) | 0.4667 | 11/11<br>(100%) | 38/52<br>(73%) | Inf<br>(0.79-Inf) | 0.1034 | 11/13<br>(85%) | 45/65<br>(69%) | 2.42<br>(0.46-24.47) | 0.3303 |
| One or more drugs tested | 1/2<br>(50%) | 10/19<br>(53%) | 0.90<br>(0.01-78.36) | 1 | 11/11<br>(100%) | 42/55<br>(76%) | Inf<br>(0.66-Inf) | 0.1036 | 12/13<br>(92%) | 52/74<br>(70%) | 5.01<br>(0.66-226.51) | 0.1701 |
| MDR | 0/2<br>(0%) | 5/19<br>(26%) | 0.00<br>(0.00-18.01) | 1 | 5/11<br>(45%) | 12/55<br>(22%) | 2.93<br>(0.60-13.93) | 0.1339 | 5/13<br>(38%) | 17/74<br>(23%) | 2.08<br>(0.47-8.38) | 0.3000 |

**Table S7.** AMR frequencies in transmitted strains compared to those not involved in transmission. Non-susceptibility was defined as I or R according to clinical breakpoints (see **Methods**). MDR was defined as non-susceptibility to ampicillin or augmentin plus at least one other antimicrobial. Associations calculated using two-sided Fisher’s exact test.

|  | *H. influenzae* |  |  |  | *H. parainfluenzae* |  |  |  | Both |  |  |  |
| --- | --- | --- | --- | --- | --- | --- | --- | --- | --- | --- | --- | --- |
|  | Transmission | Other | OR (95%CI) | *p*-value | Transmission | Other | OR (95%CI) | *p*-value | Transmission | Other | OR (95%CI) | *p*-value |
| Ampicillin | 2/3<br>(67%) | 4/17<br>(24%) | 5.79<br>(0.24-410.68) | 0.2018 | 3/3<br>(100%) | 53/73<br>(73%) | Inf<br>(0.15-Inf) | 0.5619 | 5/6<br>(83%) | 57/90<br>(63%) | 2.87<br>(0.30-140.90) | 0.4183 |
| Augmentin | 0/3<br>(0%) | 1/12<br>(8%) | 0.00<br>(0.00-155.62) | 1 | 2/3<br>(67%) | 22/70<br>(31%) | 4.27<br>(0.21-262.21) | 0.2500 | 2/6<br>(33%) | 23/82<br>(28%) | 1.28<br>(0.11-9.64) | 1 |
| Cefotaxime | 0/3<br>(0%) | 0/17<br>(0%) | 0.00<br>(0.00-Inf) | 1 | 0/3<br>(0%) | 2/72<br>(3%) | 0.00<br>(0.00-150.09) | 1 | 0/6<br>(0%) | 2/89<br>(2%) | 0.00<br>(0.00-84.49) | 1 |
| Co-trimoxazole | 1/3<br>(33%) | 7/17<br>(41%) | 0.73<br>(0.011-16.65) | 1 | 1/3<br>(33%) | 21/74<br>(28%) | 1.26<br>(0.02-25.38) | 1 | 2/6<br>(33%) | 28/91<br>(31%) | 1.12<br>(0.096-8.37) | 1 |
| Rifampicin | 0/3<br>(0%) | 2/9<br>(22%) | 0.00<br>(0.00-17.60) | 1 | 1/3<br>(33%) | 3/51<br>(6%) | 7.39<br>(0.10-186.70) | 0.2098 | 1/6<br>(17%) | 5/60<br>(8%) | 2.17<br>(0.039-26.45) | 0.4490 |
| Ampicillin or<br>Augmentin | 2/3<br>(67%) | 4/12<br>(33%) | 3.62<br>(0.15-264.27) | 0.5253 | 3/3<br>(100%) | 54/69<br>(78%) | Inf<br>(0.11-Inf) | 1 | 5/6<br>(83%) | 58/81<br>(72%) | 1.97<br>(0.20-97.76) | 1 |
| One or more<br>drugs tested | 2/3<br>(67%) | 9/17<br>(53%) | 1.73<br>(0.076-117.70) | 1 | 3/3<br>(100%) | 60/74<br>(81%) | Inf<br>(0.089-Inf) | 1 | 5/6<br>(83%) | 69/91<br>(76%) | 1.59<br>(0.16-78.76) | 1 |
| MDR | 1/3<br>(33%) | 4/17<br>(24%) | 1.58<br>(0.022-38.86) | 1 | 2/3<br>(67%) | 19/74<br>(26%) | 5.63<br>(0.28-346.75) | 0.1789 | 3/6<br>(50%) | 23/91<br>(25%) | 2.92<br>(0.37-23.35) | 0.3379 |

